# Mutability Patterns Across the Spike Glycoprotein Reveal the Diverging and Lineage-specific Evolutionary Space of SARS-CoV-2

**DOI:** 10.1101/2022.02.01.478697

**Authors:** Roberth A. Rojas Chávez, Mohammad Fili, Changze Han, Syed A. Rahman, Isaiah G. L. Bicar, Sullivan Gregory, Guiping Hu, Jishnu Das, Grant D. Brown, Hillel Haim

**Author notes:** To whom correspondence should be addressed: Hillel Haim, MD, PhD, Department of Microbiology and Immunology, The University of Iowa, 51 Newton Rd, 3-770 BSB, Iowa City, Iowa, 52242.

## Abstract

Mutations in the spike glycoprotein of SARS-CoV-2 allow the virus to probe the sequence space in search of higher-fitness states. New sublineages of SARS-CoV-2 variants-of-concern (VOCs) continuously emerge with such mutations. Interestingly, the sites of mutation in these sublineages vary between the VOCs. Whether such differences reflect the random nature of mutation appearance or distinct evolutionary spaces of spike in the VOCs is unclear. Here we show that each position of spike has a lineage-specific likelihood for mutations to appear and dominate descendent sublineages. This likelihood can be accurately estimated from the lineage-specific mutational profile of spike at a protein-wide level. The mutability environment of each position, including adjacent sites on the protein structure and neighboring sites on the network of comutability, accurately forecast changes in descendent sublineages. Mapping of imminent changes within the VOCs can contribute to the design of immunogens and therapeutics that address future forms of SARS-CoV-2.

## INTRODUCTION

Since emerging in December 2019, SARS-CoV-2 has caused devastating effects worldwide. By June 2022, more than 6 million deaths have been attributed to the infection, and estimated economic losses greater than $10 trillion are expected by the end of this year^1,2^. Several SARS-CoV-2 VOCs have appeared at different time points of the pandemic^3,4,5^. Most mutations in the VOCs that impact infection are found in the spike protein that adorns the virus surface. Spike mediates fusion with host cells and is the primary target for antibodies elicited by infection or vaccination^6^. Changes in spike can increase virus infectivity, transmissibility or resistance to vaccine-elicited antibodies and therapeutics^7^. New sublineages of the VOCs continuously appear with such mutations in spike^8,9^. Interestingly, only a minority of these changes are convergent, whereby the same substitution occurs in multiple lineages^10,11,12,13^. The latter are generally guided by positive selection pressures^14^. By contrast, most mutations that define VOC sublineages are found at distinct positions of spike and occur at evolutionarily neutral sites^15,16^. This observation raises two important questions. First, do the distinct patterns of mutations in the lineages reflect the stochastic nature of their appearance or a lineage-specific likelihood for their emergence? Second, if not driven solely by stochasticity, what clues can we identify that will inform of the evolutionary space of each lineage? Answers to these questions are critical for our ability to develop vaccines and therapeutics that can maintain their efficacy against future forms of this virus.

To address the above questions, we examined the frequency of independent mutation events at each position of spike in different SARS-CoV-2 VOCs. Lineage-specific mutational spaces were observed, as defined by the patterns of low-frequency substitutions in spike within populations infected by each lineage. We compared the mutational space of spike in each lineage with the evolutionary path of the virus within the lineage (i.e., the observed mutations that define descendant sublineages). We discovered that the sites of change in the emergent sublineages were characterized by high mutability in the ancestral lineage and a high mutability “environment”, composed of adjacent positions on the protein structure and co-mutable sites across the protein. These measures of positional and environmental variability predicted remarkably well the changes observed in the new sublineages of the VOCs. Our studies of spike mutability at the protein-wide level reveal the diversifying nature of the evolutionary space of this protein and demonstrate the high predictability of the changes that give rise to new SARS-CoV-2 spike forms.

## RESULTS

### The mutational space of SARS-CoV-2 spike is lineage-specific and diversifying

We examined the sites of mutation in the spike protein that define sublineages of SARS-CoV-2 VOCs. Using designations of the Pango classification system^17^, the lineages compared were B.1.1.7 (VOC Alpha), P.1 (Gamma), AY.4 (Delta), and lineages BA.1 and BA.2 (Omicron). Sites of mutation in descendant sublineages of the above that emerged until April 8^th^ 2022 were compared. In addition, we calculated for these sites the synonymous and nonsynonymous mutation rates in each lineage to infer the sites under positive selection. As expected, most sites of change that define descendent sublineages did not show evidence for positive selection in their ancestral lineages (**Fig. 1a**). Importantly, limited similarity was observed between the sites of mutations in the different VOCs – only three of the 41 sublineage-defining mutation sites appeared in more than one VOC.

**Figure 1.**
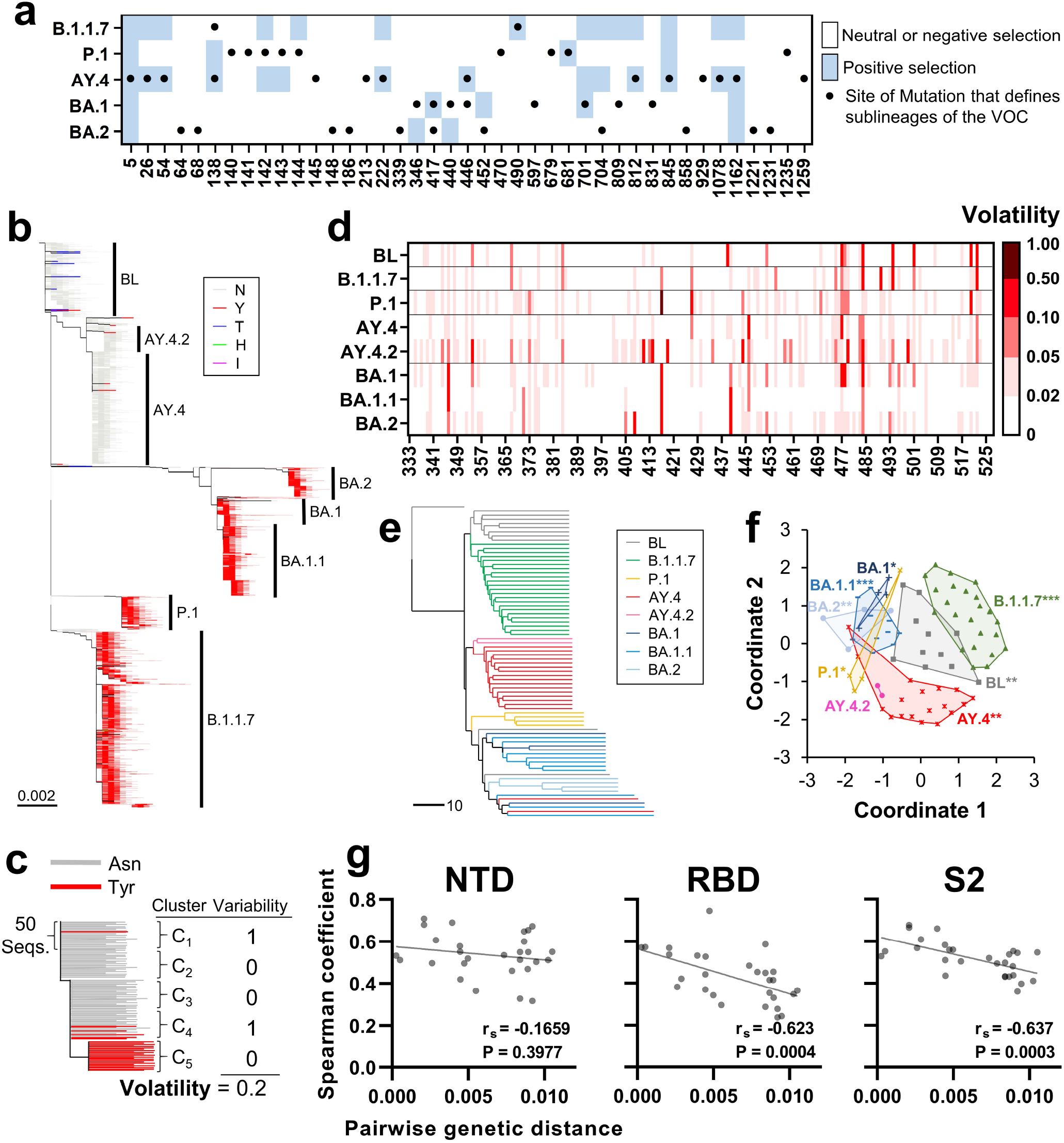
The mutational space of the spike protein is specific for SARS-CoV-2 lineage. **(a)** Sites of mutation that define sublineages of the indicated VOCs are shown by black dots. Shaded cells indicate sites under positive selection pressure in the VOCs. **(b)** Phylogenetic tree based on 40,350 unique spike sequences from the indicated lineages. The baseline (BL) group is defined as sequences within a distance of 0.0015 nucleotide substitutions per site from the SARS-CoV-2 ancestral strain, **(c)** Schematic of our approach to calculate volatility for each position of spike, **(d)** Volatility calculated at RBD positions using sequences from the indicated VOCs or the baseline group, **(e)** Each lineage was partitioned into clusters of 500 sequences. The absence or presence of volatility in each cluster was determined, and each assigned a 1273-bit string that describes the volatility at all spike positions. Strings were compared using the UPGMA clustering method, (f) Euclidean distances between the strings were calculated and the centroid of each lineage determined. A permutation test was used to compare between the mean in-lineage versus between-lineage distances. P-values: *, P<0.05; **, P<0.005; ***, P<0.0005. **(g)** Volatility was calculated in each lineage at all positions of the NTD (20-286), RBD (333-527) and S2 (686-1213). The Spearman correlation coefficient between volatility values in any two lineages was calculated. Coefficients are compared with the mean genetic distance that separates any two lineages. r_s_, Spearman coefficient. P-values, two-tailed test.

To investigate the basis for these distinct patterns of change, we first compared the mutational space of spike between the above lineages, as defined by the collection of sites that exhibit amino acid variability among strains that are phylogenetically closest to the lineage ancestor. In addition, as examples, we included the highly-prevalent sublineages BA.1.1 and AY4.2, which emerged from lineages BA.1 and AY.4, respectively. All other Pango-designated sublineages of the above variants were removed from the datasets. As a group representative of isolates closest to the SARS-CoV-2 ancestral strain (designated the SARS-CoV-2 “baseline”), we used sequences within 0.0015 substitutions per site from the SARS-COV-2 spike ancestral sequence (Wuhan strain, accession number NC_045512)^18^. Spike sequences that appeared at least twice in the population were aligned and “compressed” to obtain a single representative for each unique sequence. Evolutionary relationships among them were inferred, and a maximum likelihood phylogenetic tree was constructed (see Methods and **Fig. 1b**). To quantify amino acid variability at each position of spike, the lineages were partitioned into clusters of 50 sequences based on phylogeny, and the proportion of clusters that contain amino acid variability at each position was calculated (**Fig. 1c**). We designate this measure of variability “volatility”, which quantifies the frequency of substitution events.

Considerable differences were observed between the volatility profiles of spike in the diverse lineages (see RBD positions in **Fig. 1d** and all spike positions in **Supplementary Fig. 1a** and **1b**). To determine the relationships between volatility patterns, we partitioned each lineage into groups of 10 clusters. The absence or presence of amino acid variability at each spike position was determined for each group, and all groups were assigned 1273-bit vectors that describe their volatility profiles at all spike positions. Jaccard distances between the vectors were applied to determine hierarchical relationships using the unweighted pair group method with arithmetic mean (UPGMA) approach^19,20^. Clear clustering of group profiles from the same lineage was observed (**Fig. 1e**). To determine statistical significance of these patterns, Euclidean distances were measured between all vectors, and intra-lineage distances compared with inter-lineage distances using a permutation test^21^. As shown in **Fig. 1f**, all lineages (except the smaller AY4.2) exhibited significant specificity of their volatility patterns.

To quantify divergence of the volatility patterns, we performed pairwise comparisons between the volatility profiles of any two lineages and the genetic distance that separates them. Positions of the RBD, N-terminal domain of spike (NTD) and S2 subunit were analyzed separately. The correlation coefficient for any lineage pair was then compared with the nucleotide distance between the lineage founders. For the RBD and S2 subunits, a strong negative relationship was observed between the genetic distance that separates any two lineage founders and the correlation between their volatility profiles (**Fig. 1e**). For the NTD, which contains a relatively high proportion of sites with mutations, we did not observe such a relationship.

These findings suggest that the mutational space of spike (i.e., the collection of sequence states that are sampled) is specific for each lineage and is diversifying. As such, and assuming that the mutational space corresponds with the evolutionary path of the virus in the population, these findings also suggest that each lineage may have a distinct set of changes that can appear in descendent sublineages. This possibility was explored in the studies described below.

### A high volatility state and a high volatility environment increase the likelihood of spike positions to emerge with founder mutations in descendent lineages

We examined the relationship between the volatility of any spike position in a population of related strains and the emergence of mutations at this position in descendent lineages. To this end, we first analyzed mutability profiles in the SARS-CoV-2 baseline that preceded emergence of the VOCs. These profiles were compared with the mutations that define the emergent lineages. For simplicity, we focused these analyses on all 615,374 spike sequences from samples collected worldwide between December 2019 and July 2021. Evolutionary relationships among the nucleotide sequences were inferred and a maximum likelihood tree was constructed (**Fig. 2a**). We then partitioned the tree into discrete groups separated by a minimal distance of 0.004 substitutions per site. As expected, many groups corresponded to known SARS-CoV-2 VOCs. The baseline groups were distinguished from the terminal emergent groups (G_T1_-G_T8_) using a threshold of 0.0015 substitutions per site between the centroid of each group and the SARS-CoV-2 spike ancestral sequence. All groups are described in **Supplementary Table 1**.

**Figure 2.**
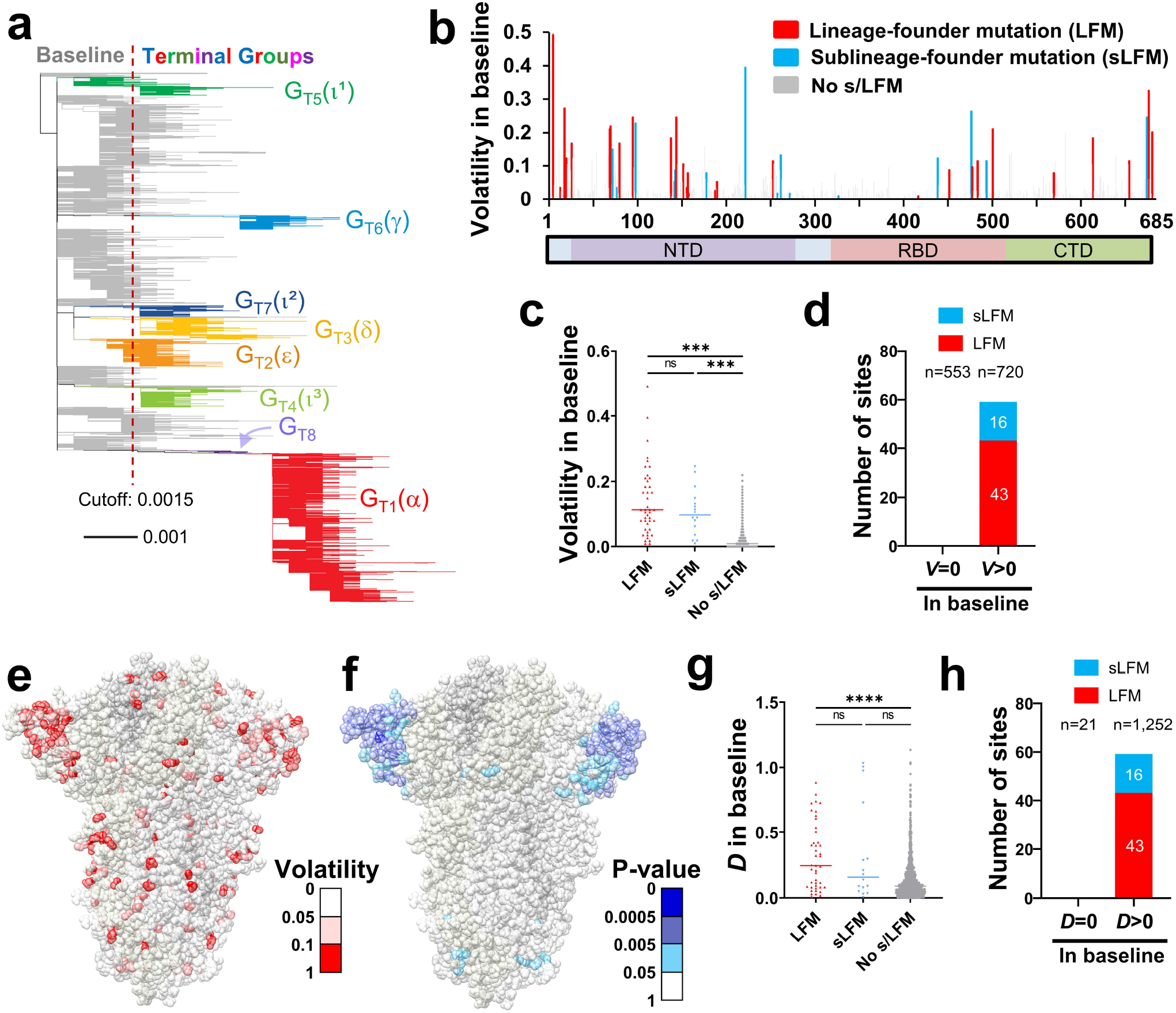
Spike positions with high volatility or a volatile environment emerge as sites of lineage-founder mutations. **(a)** Phylogenetic tree based on 16,808 unique spike sequences. Terminal groups are colored and labeled, with their WHO variant designations in parentheses, **(b)** Volatility values for all positions of spike subunit S1 calculated using the 114 baseline clusters (see values for subunit S2 in **Supplementary Fig. 2b). (c)** Comparison of volatility values between spike positions that emerged with LFMs, sLFMs or no such mutations. P-values in an unpaired T test: ***, P<0.0005; ****, P<0.00005; ns, not significant, **(d)** Number of positions that appeared with LFMs and sLFMs when volatility (V) in the baseline group was zero or larger than zero. The number of positions in each subset (n) is indicated, **(e)** Mapping of the volatility values calculated in the baseline group onto the spike trimer (PDB ID 6ZGI). **(f)** Results of a permutation test to identify positions with high volatility at their 10 closest neighbors on the trimer structure. Low P-values indicate a high-volatility environment, **(g)** The variable *D* describes for each position the total distance-weighted volatility at adjacent positions on the spike trimer. *D* values are compared between positions with LFMs, sLFMs or no such mutations, **(h)** The number of positions that emerged with LFMs or sLFMs when the *D* value was zero or larger than zero.

Volatility at each spike position in the baseline was compared with the absence or presence of two types of mutations: **(i) Lineage-founder mutations (LFMs)**, which are found in the group ancestors and in at least 50% of all sequences from that group, and **(ii) Sublineage-founder mutations (sLFMs)**, which are not found in the group ancestor and represent clonal expansions that dominate at least one 50-sequence cluster but less than 50% of all group clusters (see examples in **Supplementary Fig. 2a**). A total of 43 LFMs and 16 sLFMs were detected in the baseline and terminal groups (see **Supplementary Table 1**). Most positions with high volatility in the baseline emerged with LFMs or sLFMs (see positions of spike subunit S1 in **Fig. 2b** and of subunit S2 in **Supplementary Fig. 2b**). Among positions with the highest volatilities, most appeared as s/LFMs in at least one group (**Supplementary Fig. 2c**). Sites of s/LFMs were more volatile than sites with no such mutations (**Fig. 2c**). Furthermore, non-volatile sites in the baseline did not emerge with s/LFMs in any baseline or terminal group (**Fig. 2d**). Therefore, for any given site, a high level of volatility (i.e., a high frequency of independent mutation events) in the baseline group precedes (as inferred phylogenetically) the emergence of s/LFMs in the descendent lineages.

We recently examined the within-host patterns of amino acid variability in the envelope glycoproteins (Envs) of human immunodeficiency virus type 1 (HIV-1)^22^. We found that the variability at many positions of the CD4-binding site can be accurately estimated by the variability at adjacent positions on the three-dimensional structure of the protein. Analysis of the spatial distribution patterns of volatile sites on the SARS-CoV-2 spike structure suggested a similar clustering of volatility at multiple loci, most notably in the NTD (see **Fig. 2e** and sites with statistically significant clustering in **Fig. 2f**). We hypothesized that if such associations are “stable” over time, then the likelihood for future changes at any position may be associated with volatility of its neighboring positions. To test this hypothesis, we generated a variable (designated *D*) that describes for each position *i* the total distance-weighted “environmental” volatility:

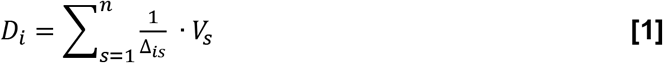

where *n* is the number of positions s within 6 Å of position *i, Δ_is_* is the distance between the closest two atoms of positions *i* and each position s, and *V_i_* is the volatility at each position s. Similar to the volatility values, *D* values were higher for positions that emerged with s/LFMs (**Fig. 2g**). Furthermore, positions with a non-volatile environment (i.e., a *D* value of zero) did not emerge with s/LFMs (**Fig. 2h**). Therefore, high volatility at any position in the SARS-CoV-2 baseline and high volatility at adjacent positions on the protein increase the likelihood of the site to emerge with s/LFMs in descendent lineages.

### Volatility at neighboring sites on the network of co-volatility increases the likelihood of spike positions to emerge with lineage founder mutations

We examined whether the clustering patterns of volatility at adjacent positions on spike can be generalized to describe associations that are not dependent on physical proximity. To this end, we used the 114 baseline clusters to determine the co-occurrence of volatility at any two spike positions within the clusters (see schematic in **Fig. 3a**). P-values were calculated using Fisher’s exact test and used to construct the network of co-volatile sites, whereby the edges that connect the nodes (positions) are defined by the statistical significance of the association between their volatility patterns (see example of a network segment in **Fig. 3b** and distribution of P-values in **Supplementary Fig. 3a**). To determine the significance threshold to apply for network construction, we examined structural properties of the network and its robustness to random deletion of edges. Two network topological metrics were assessed: **(i)** Degree distribution, which describes the average number of connections each node has with other nodes, and **(ii)** Closeness centrality, which describes for each node the sum of the path lengths to all other nodes in the network (more central nodes have lower values)^23^. For robust scale-free networks, limited random-edge deletions only minimally perturb their topological properties^24^. We found that networks defined at a more stringent significance threshold (P<0.01) were more robust to edge deletions, with minimal impact on both degree distribution and closeness centrality at the expense of losing information (**Fig. 3c** and **Supplementary Fig. 3b**). By contrast, when less stringent significance thresholds were used (P<0.1), the number of edges was greater (i.e., they contained more information regarding the co-volatile positions); however, the network was less robust to edge deletions. This suggested that an intermediate significance threshold (P<0.05) would provide a sufficiently stable network without losing most information.

**Figure 3.**
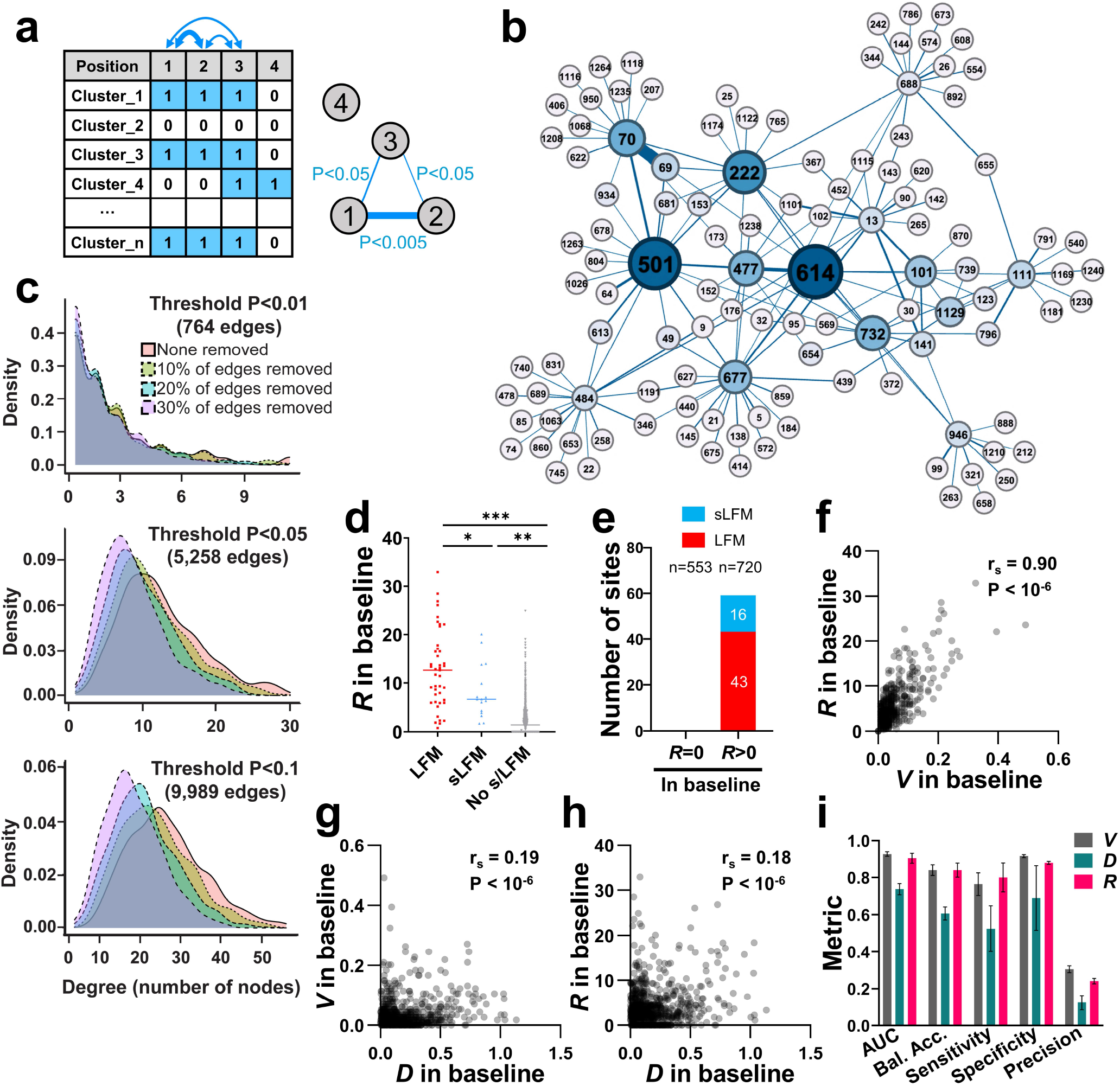
High volatility at co-mutable sites is associated with emergence of LFMs and sLFMs. **(a)** Schematic of our approach to calculate co-volatility of spike positions. For all positions, the absence (0) or presence (1) of amino acid variability was determined in each cluster of 50 sequences. The co-occurrence of a volatile state at all position pairs was determined using Fisher’s test, and the P-values were used to construct the network of co-volatility between all positions, (b) The co-volatility network around position 614 as the root node. Edges were assigned to positions pairs if the P-value was smaller than 0.05. First- and second-degree nodes are shown. Node size corresponds to the number of triangle counts for each position, **(c)** Network robustness analyses. Networks were constructed using P-value thresholds of <0.01, <0.05 or <0.1. For each network, we randomly deleted 10%, 20% or 30% of edges and examined the effect on network stability. The degree distribution (i.e., the number of nodes associated with each position) is shown for the intact and depleted networks, **(d)** *R* values, which describe for each position the total weighted volatility at network-associated positions was calculated. *R* values for spike positions that emerged with LFMs, sLFMs or with no such mutations are shown, **(e)** Number of LFMs and sLFMs that emerged at spike positions when *R* in the baseline was equal to zero or greater than zero, **(f-h)** Correlations between V, *D* and *R* values calculated using the baseline sequences. r_s_, Spearman coefficient. P-value, two-tailed test, **(i)** Classification metrics for evaluating performance of *V, D* and *R* to predict presence of s/LFMs. Error bars, standard errors of the means for five-fold cross validation.

We examined whether, for any position *i* of spike, presence of high volatility at its network-associated co-volatile sites (q) is associated with emergence of s/LFMs. To this end, we generated a simple measure (*R*) designed to capture for each position *i* the total volatility of its network-neighbors q (q_1_ q_2_, q_3_ … q_n_), using a P-value of 0.05 as the threshold:

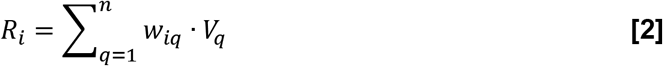

where *V*_q_ is the volatility at each position q calculated using the baseline sequences, and *w*_iq_ is the evidence for association between volatility of position *i* and each of its positions q (calculated as the -log_10_(P-value) in Fisher’s test). Similar to the *V* and *D* values, *R* values were significantly higher for positions with s/LFMs relative to positions with no such mutations (**Fig. 3d**). Furthermore, an *R* value of zero in the baseline was invariably associated with lack of s/LFM appearance (**Fig. 3e**). Overall, *V* and *R* values for any position correlated well, and considerably better than their correlation with *D* (**Fig. 3, f-h**). Nevertheless, as shown below, *V* and *R* values exhibit different levels of predictive performance when small population sizes are tested.

We compared the performance of the three variables (*V, D* and *R*) to predict the emergence of LFMs or sLFMs using a univariate logistic regression model. Higher classification metrics were observed for *V* and *R* relative to *D*, with area under the receiver operating characteristic curve (AUC) values higher than 0.9 for both *V* and *R* (**Fig. 3i**). In comparison, precision of these variables was modest, at 0.3 for *V* and 0.24 for *R*, indicating a relatively high false-positive rate. Taken together, these findings show that the emergence of s/LFMs at any spike position is associated with a state of high volatility in the ancestral lineage, as well as high volatility at adjacent positions on the protein and at associated sites on the co-volatility network.

### Volatility profiles among sequences from the early pandemic capture the mutational patterns of the emergent lineages

We examined whether a combination of the volatility-based variables would better capture the observed evolutionary path of the virus than each of them separately. To this end, we indexed all sequences by the time of sample collection and tested whether viruses that temporally preceded emergence of SARS-CoV-2 lineages can predict the mutations they contain. For these analyses, sequences were classified by their Pango lineage designations rather than our phylogeny-based group definitions. We first determined the formation time of each lineage, defined as the date by which 26 unique sequences from the lineage were detected (see **Fig. 4a** and **Supplementary Table 2**). Based on these timelines, we divided the sequences into an “early-phase” group that is used to predict emergence of the LFMs in the “lineage-emerging phase”. The early-phase group included one sequence from lineage B.1.1.7 and none from the other emergent lineages. Six minor lineages emerged early in the pandemic that contained mutations at positions 614, 222 and 477 (see **Supplementary Table 3**). To avoid a potential bias, these positions were excluded from our analyses. A total of 67 LFM sites were identified in the lineage-emerging phase.

**Figure 4.**
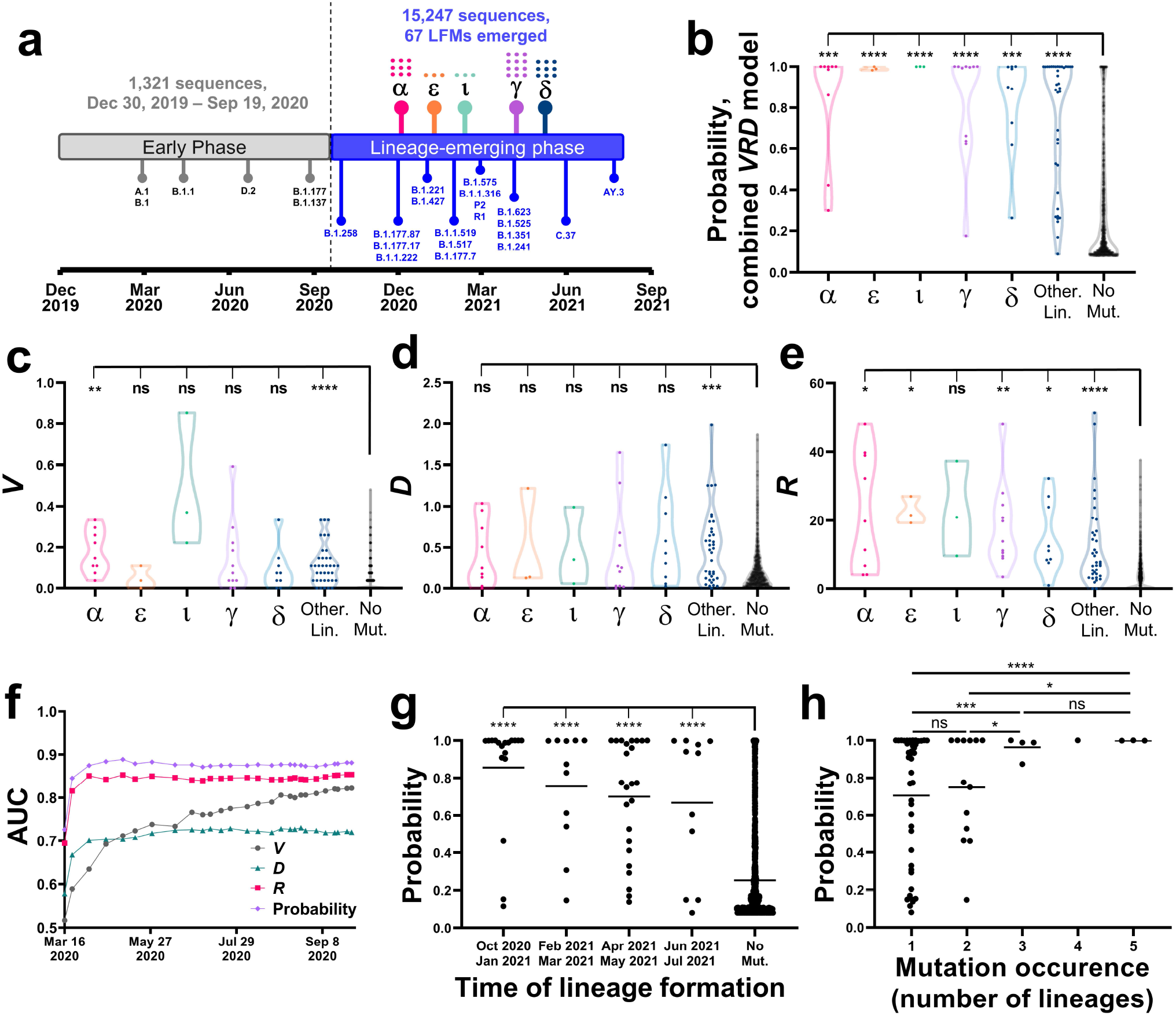
Volatility patterns among early-pandemic viruses predict emergence of mutations during the lineage-emerging phase. **(a)** Timeline for emergence of SARS-CoV-2 lineages until July 2021. Lineage emergence time is defined as the date by which 26 sequences that contain all lineage mutations were identified. Lineages with WHO variant designations are indicated by their symbols (see list in Supplementary **Table 2**) and the number of LFMs in each is shown by dots, **(b)** *V, D* and *R* values were calculated for all spike positions using the early phase sequences and applied to a logistic regression model to predict emergence of LFMs. Datapoints describe probabilities assigned to all spike positions and are grouped by the lineage in which they emerged as LFMs. Values are compared between the LFM sites in the indicated VOCs (or minor lineages, labeled “Other Lin.”) and the no-mutation (“No mut”) sites. Significance of the differences was calculated using an unpaired T test: *, P<0.05; **, P<0.005; ***, P<0.0005; ****, P<0.00005; ns, not significant, **(c-e)** V, *D* or *R* values calculated for all spike positions using the early phase sequences are grouped as described in panel b. **(f)** *V, D* and *R* values and the combined probability were calculated using sequences from different time points of the early phase. AUC values are shown for each time point, for predicting emergence of the 67 LFMs in the lineage-emerging phase. The number of spike sequences at each time point is indicated, **(g)** LFM sites were grouped by the emergence time of the first lineage that contains them. Mutation probabilities assigned to the sites by sequences collected until April 1^st^ 2020 are compared with the probabilities assigned to the no-mutation sites, **(h)** Probabilities assigned by the April 1^st^ 2020 dataset are shown for LFM sites that appeared in one or more lineages. Values are compared between all groups using an unpaired T test.

The early-phase sequences were divided into 27 clusters of 50 sequences, which were used to calculate *V, D* and *R* values. These values were applied to a logistic regression model that was trained to predict the emergence of LFMs using the phylogeny-indexed baseline sequences (see Methods section). The output of the model is the probability of each site to emerge with LFMs in the lineage-emerging phase. For all VOCs tested, as well the non-VOC lineages (analyzed collectively), the probabilities calculated for LFM sites were significantly higher than probabilities assigned to the non-LFM sites (**Fig. 4b**). Predictions based on the combined model exhibited considerably higher performance than those based on the individual variables (**Fig. 4, c-e**).

To examine the changes in probabilities assigned to the sites of mutation during the early stages of the pandemic, we calculated *V, D* and *R* values and the combined probability using increasing numbers of sequences indexed by the time of sample collection. Interestingly, the pattern of LFMs was predicted with high sensitivity and specificity by the time three clusters were formed (150 unique sequences), corresponding to samples collected until April 1^st^, 2020 (**Fig. 4f** and **Supplementary Fig. 4, a-c**). Of the individual predictors, *R* exhibited the highest performance, modestly lower than the combined probability, whereas performance of *V* gradually increased. Further analysis of the performance of the first three clusters showed that higher probabilities were assigned to mutation sites of lineages that emerged at earlier stages of the pandemic (**Fig. 4g** and **Supplementary Fig. 4d**). Higher probabilities were also assigned to convergent sites (i.e., those that emerged with LFMs in multiple lineages, **Fig. 4h** and **Supplementary Fig. 4e**).

We note that while *V* and *R* values calculated using all sequences of the baseline group correlated well (**Fig. 3f**), the performance of *R* was considerably higher when a small number of sequences was available (**Fig. 4f**). For example, analysis of the two major VOCs circulating during the lineage-emerging phase showed near-maximal *R* values for most LFM sites in March 2020 whereas *V* values of these sites gradually increased over time (**Supplementary Fig. 5**).

Taken together, these findings show that a high level of volatility at any site and at its spatial- and network-associated sites precedes emergence of mutations in descendant lineages. A small number of sequences is required to accurately estimate the likelihood of sites for emergence as LFMs. Total volatility at network-associated sites exhibits a higher level of sensitivity at earlier stages of the pandemic than volatility values of the sites.

### Spike mutations in emerging SARS-CoV-2 sublineages are accurately forecasted by patterns of volatility in their ancestral lineages

We examined the patterns of volatility among sequences that preceded emergence of the VOC sublineages. For this purpose, we focused on SARS-CoV-2 lineages from the major VOCs, including B.1.1.7, P1, AY.4, BA.1 and BA.2. Sequences from the baseline of each lineage were used to forecast the mutations that define its descendent sublineages. All emergent sublineages with Pango designations and all clusters of 50 sequences that contain a non-lineage-ancestral residue as the majority variant at any site were excluded from the datasets. The remaining clusters were used to calculate *V, D* and *R* values and to assign a mutation probability to each position. Two mutation types were tested as outcomes: **(i)** Mutations that define Pango sublineages of the VOCs, and **(ii)** Mutations that are dominant in two or more 50-sequence clusters of the lineage (but are not assigned a Pango sublineage designation). As shown in **Fig. 5a**, both outcomes were predicted well using the baseline sequences of each variant (see **Supplementary Table 3** for probability values). To determine the lineage specificity of the predictions, we compared the probabilities assigned to all sites of each lineage with the mutational outcomes in all other lineages. Consistent with the above findings, the highest AUC values were observed for predictions of the changes that occurred in the homologous lineage (**Fig. 5b**).

**Figure 5.**
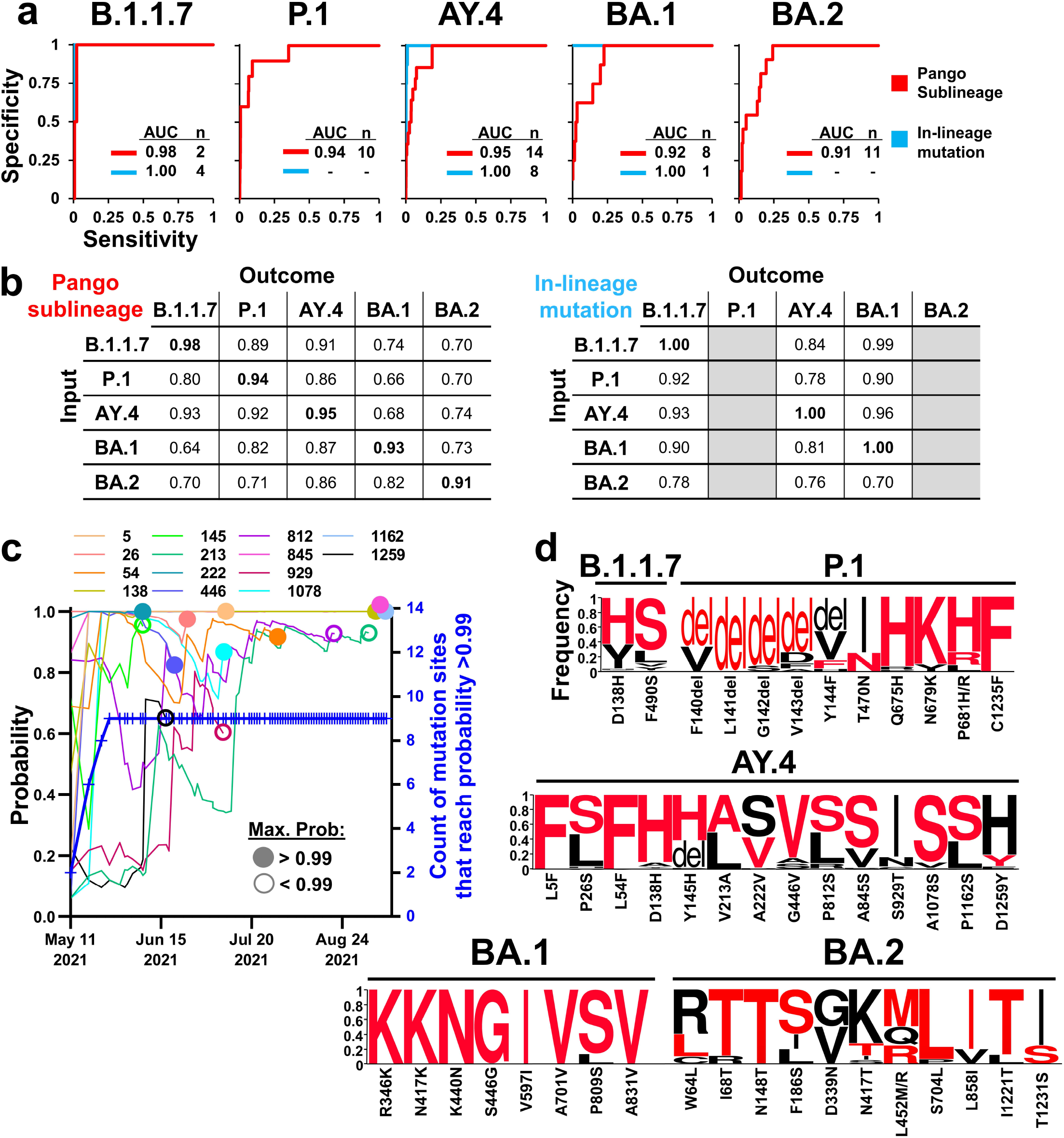
The mutational patterns of new SARS-CoV-2 sublineages are accurately captured by volatility profiles in their ancestral lineages. **(a)** Sequences from the baseline of the indicated VOCs were used to calculate the combined probability for mutations at all spike positions. Values were compared with two outcomes: (i) Emergence of a mutation defines a new Pango sublineage (in red), or (ii) Presence within the lineage baseline of two or more 50-sequence clusters with a dominant non-lineage-ancestral residue (in blue). The number of events detected for each outcome (n) and the AUC values are shown, **(b)** Mutation probabilities calculated using the sequences of each lineage are compared with the outcomes observed in all lineages, **(c)** AY.4 sequences were indexed by sample collection date. The cumulative probabilities for the 14 sites that emerged as sublineages of AY.4 are shown. Circles indicate the time of sublineage emergence. Filled circles indicate that the site reached a probability of 0.99. The blue line indicates for each time point the number of sites that reached a probability of 0.99 **(d)** Weblogos describe the frequencies of minority variants at sites that appeared in the VOC sublineages by April 8^th^ 2022. Frequencies are expressed as a percent of all sequences with a non-lineage-ancestral residue. The residues that emerged in the sublineages are shown in red font. The residue change in each lineage from the SARS-CoV-2 ancestor is shown below the axis.

We also investigated the changes in probabilities assigned to the sites of sublineage mutations from the time their ancestral lineage had emerged. For these tests, we focused on AY.4, which has circulated in the population for a longer time period than other lineages (global emergence in May 2021). Mutation probabilities were calculated using increasing amounts of sequences indexed by time (**Fig. 5c**). Similar to the mutations that define lineages B.1.1.7 and B.1.617.2 (**Supplementary Fig. 5**), most sites of change in the sublineages of AY.4 exhibited high mutation probabilities at early stages after AY.4 emergence. Of the 14 sites of mutation in AY.4 sublineages, nine surpassed the 0.99 probability threshold (mean 95^th^ probability percentile) at least once during the first month after emergence of AY.4. We note that in these tests a positive outcome was defined as a mutation that appeared in immediately descendent sublineages (e.g., mutations that define AY.4.2 but not mutations that define AY.4.2.1). Nevertheless, several sites with high mutation probabilities were also observed in second-order lineages. For example, the second-highest probability in BA.1 was assigned to position 1081. Since this change was observed in the second-order sublineage BA.1.15.1, as designated by the Pango system, it was not considered a positive outcome site.

Finally, we examined the specific changes in amino acid occupancy at the sites of mutation in the emergent sublineages. In most cases, the minority variant with the highest frequency in the baseline group of each lineage also appeared as the dominant residue in the new sublineage (see characters in red font in **Fig. 5d**). These findings further support the notion that the evolutionary path of spike within each lineage is accurately captured by its mutational space across all sites of the protein, as detected among early isolates of the lineage.

## DISCUSSION

Since January 2022, SARS-CoV-2 variant Omicron has dominated the landscape of VOCs circulating worldwide. Both major lineages of Omicron (BA. 1 and BA.2) contain mutations in spike that reduce virus sensitivity to immune sera and COVID-19 therapeutics^25,26^. To address these unique antigenic properties^27,28^, Omicron-specific immunogens have been developed and tested^29,30,31^. Nevertheless, new sublineages of this VOC emerge with mutations in spike that further impact virus transmissibility and sensitivity to COVID-19 vaccines^8,9^. Most changes that give rise to new sublineages do not appear to be driven by positive selective pressures (**Fig. 1a**). Instead, they occur at evolutionarily neutral positions and “hitchhike” onto the driver mutations^32^. As such, it would be expected that the evolutionary space available for spike (i.e., the sequence states that can be occupied by expanding lineages of the virus) would be large and driven by the stochastic nature of the hitchhiking event. Here we introduce a simple probabilistic definition of the evolutionary space of the virus. We show that, in fact, diverse SARS-CoV-2 lineages have vastly distinct evolutionary spaces. Each position of spike has a specific and measurable likelihood to appear as a dominant mutation within descendants of each lineage.

The volatility of each site only partially captures its likelihood for emergence with LFMs. More accurate estimates are provided by the mutability profile of spike at the whole-protein level, including adjacent sites on the protein and co-volatile sites. The latter variable is more sensitive to the changes at shorter time frames from emergence of the parental lineage. How does volatility of the “environment” capture the mutability of each site? Clustering of volatile sites on the linear sequence of the protein can be explained by mutational hotspots due to properties of the viral RNA^33,34^. Clustering on the three-dimensional structure can be explained by high permissiveness of the region for change due to their limited impact on fitness^35^. By contrast, the association between volatility of sites separated by larger distances on the protein is less intuitive. We propose that such associations describe the epistasis network of spike (i.e., the sites that the amino acid occupancy of one affects the fitness of another). Indeed, the volatility of each position likely captures its fitness profile; low volatility describes a state with a single high-fitness residue, whereas high volatility describes the presence of multiple residues with high fitness. Accordingly, we propose that co-volatility patterns may capture the associations between the fitness profiles of the sites. Comparison of co-volatility network structure with structure of the epistasis network of spike will address this question, and may allow us to identify the adaptive sites required to facilitate changes at sites that negatively affect virus fitness^35,36^.

We note that, despite the high predictive performance shown, these studies constitute a relatively simple framework to understand specific factors associated with the changes in SARS-CoV-2 spike. A more complete understanding will be generated by incorporating additional factors, including selective pressures applied on each site, *in vitro* fitness profiles^35^, and possibly patterns of mutations within the host. Furthermore, from a computational perspective, our strategy can be refined by applying alternative methods to define the architecture of the co-volatility network and by using more sophisticated learners to combine the volatility-based variables. In these studies, we have chosen to apply a simple logistic regression model to demonstrate the predictable nature of the changes. Importantly, the use of more homogenous donor populations, divided by their infection and vaccination status, will allow us to account for the effects of the immune response on the evolutionary path of each variant.

The predictable nature of the changes in the spike protein suggest that immunogens and therapeutics can be designed to effectively address future forms expected to dominate in the population. Advance notice of the imminent changes in each lineage allows testing of their impact on virus fitness and sensitivity to immune sera^37^. Knowledge of the sites that are not expected to change is equally important. For example, several mutations in the RBD that affect virus sensitivity to antibodies, including L452R and T478K were assigned high probabilities to occur from the baseline group, but low probabilities to occur within lineage B.1.1.7. Similarly, the convergent N501Y mutation^10^ was assigned a high probability by the baseline group but a low probability in AY.4 (data not shown). Accordingly, such mutations were not observed in sublineages of the above variants. These findings further support the notion that immunogens should be tailored to the evolutionary space that is specific to each lineages, which can be inferred at early stages after it emerges in the population.

## METHODS

### Sequence alignment

Nucleotide sequences of SARS-CoV-2 isolated from humans were downloaded from the National Center for Biotechnology Information (NCBI) database, the Virus Pathogen Database and Analysis Resource (ViPR) and from the GISAID repository^38^. The following processing steps and analyses were performed within the Galaxy web platform^39^. First, excess bases were trimmed using Cutadapt, using 5’-ATGTTTGTT-3’ and 3’-TACACATAA-5 “adapters” that flank the spike gene. Adapter sequences were allowed to match once with a minimum overlap of 5 bases, an error rate of 0.2 with a sequence length between 3,700 and 3,900 bases. All sequences with any nucleotide ambiguities were removed by replacing the non-standard bases with ‘N’ using snippy-clean_full_aln, followed by filtration of N-containing sequences using Filter FASTA. Sequences that cause frameshift mutations were excluded using Transeq. Nucleotide sequences were aligned by MAFFT, using the FFT-NS-2 method^40^. The aligned sequences were then “compressed” using Unique.seqs to obtain a single representative for each unique nucleotide sequence^41^. Nucleotide sequences were then translated with Transeq and aligned with MAFFT, FFT-NS-2^40^. The first position of each PNGS motif triplet (Asn-X-Ser/Thr, where X is any amino acid except Pro) was assigned a distinct identifier from Asn. All phylogenetic analyses were performed using the full-length spike protein, which include several sequences with amino acid insertions. To maintain consistent numbering of spike positions, all calculations described in this work were performed for the 1,273 positions of the spike protein in the SARS-CoV-2 reference strain (accession number NC_045512).

### Phylogenetic tree construction and analyses

A maximum-likelihood tree was constructed for the aligned compressed nucleotide sequences using the generalized time-reversible model with CAT approximation (GTR-CAT) nucleotide evolution model with FASTTREE ^42^. The tree was rooted to the sequence of the SARS-CoV-2 reference strain with MEGAX^43^. To divide the tree into “Groups” of sequences, we used an in-house code in Python (see link to GitHub repository in the Data Availability section). This tool uses the Newick file to divide the dataset into sequence groups with a user-defined genetic distance between their centroids. For all analyses we used a distance of 0.004 nucleotide substitutions per site for group partitioning. Groups that did not contain at least 50 unique sequences were excluded. To discern between baseline groups and terminal groups, we used a distance of 0.0015 nucleotide substitutions per site between each group centroid and the SARS-CoV-2 reference strain.

### Calculations of volatility

To calculate volatility of spike positions, we divided all sequences of each group into clusters of 50 sequences. Sequence variability in each cluster was quantified using two approaches. To calculate volatility *(V)* values, we used a binary approach, whereby each position in a 50-sequence cluster was assigned a value of 1 if it contains any diversity in amino acid sequence, or a value of 0 if all sequences in the cluster contain the same amino acid. Thus, each cluster is assigned a 1,273-feature vector that describes the absence or presence of volatility at each position of spike. Volatility was then calculated by averaging values by position across all clusters. For calculations of *D* or *R* values for each position *i,* we used a quantitative approach to define volatility at positions associated with *i* (i.e., at positions s and q in **Equation 1** and **Equation 2**, respectively). Briefly, sequence variability within each cluster was measured by assigning distinct hydropathy scores to each amino acid according to a modified Black and Mould scale^44^. The Asn residue of PNGS motifs and deletions were also assigned unique values. The values assigned were: PNGS, 0; Arg, 0.167; Asp, 0.19; Glu, 0.203; His, 0.304; Asn, 0.363; Gln, 0.376; Lys, 0.403; Ser, 0.466; Thr, 0.542; Gly, 0.584; Ala, 0.68; Cys, 0.733; Pro, 0.759; Met, 0.782; Val, 0.854; Trp, 0.898; Tyr, 0.9; Leu, 0.953; Ile, 0.958; Phe, 1; deletion site, 1.5. Variability in each cluster was calculated as the standard deviation in hydropathy values among the 50 sequences, and variability values of all clusters were averaged to obtain the volatility value for each position s or q (i.e., *V*_s_ or *V*_q_, respectively).

### Lineage specificity of volatility patterns

To determine relationships between volatility profiles of spike in the diverse lineages, we partitioned each lineage into 10-cluster groups (500 sequences). All sublineages with Pango designations and all 50-sequence clusters with a dominant non-lineage-ancestral residue at any spike position were removed from the datasets. Within each group, the absence or presence of amino acid variability at each spike position was determined. All groups were thus assigned 1273-bit strings that describes the absence (0) or presence (1) of volatility at each position of the protein. Jaccard distances between the strings were calculated and all groups compared using the Unweighted Pair Group Method with Arithmetic Mean (UPGMA) method^19^,^20^. The output (in Newick format) was used to generate a dendrogram plot with MEGAX.

To determine the lineage specificity of the volatility patterns, we used a modification of an approach we previously described^21^. Briefly, each SARS-CoV-2 lineage was divided into groups of 10 clusters (500 sequences). Volatility in the groups was calculated for all positions of spike, and each group assigned a 1273-feature vector that describes the level of volatility at all positions of spike. To compare the vectors, we first calculated for each lineage *L* the coordinates of the centroid (*C_L_*) among vectors from the same lineage. The mean intra-lineage distance (*d_intraiineage_*) was calculated as the average Euclidean distance between the lineage centroid *C_L_* and all groups from the same lineage *G_L_*, formally 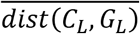. In addition, we calculated the mean inter-lineage distance (*d_interlineage_*) as the average Euclidean distance between the centroid of lineage L and all other lineage centroids 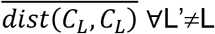. We define the ratio as:

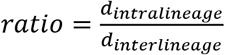. The baseline ratio (S_base_) was calculated as the *ratio* using the non-permuted data. Under the null assumption concerning the evolution of volatility profiles, the intra-lineage distances are expected to be comparable to the inter-lineage distances, while under the lineage-specific alternative, we expect clustering of volatility profiles within each lineage even across distinct 10-cluster groups. To test this, lineage identifiers were permuted and randomly assigned to each group, from which the permuted ratio (S_rand_) was calculated. The permutation process was repeated 10,000 times. The P value was calculated as the fraction of the 10,000 tests that S_rand_ was smaller than S_base_.

### Co-volatility network construction

To determine the co-volatility of any two spike positions, we generated a matrix that contains binary volatility values in all clusters of the tested group (rows) for all 1,273 spike positions (columns). The co-occurrence of a volatile state between any two spike positions was calculated using Fisher’s exact test and the associated P-value determined using a custom Java script. To construct the network of co-volatility, we used as input the matrix that describes the - log_10_(P-value) between the volatility profiles of any two spike positions, whereby nodes are the positions of spike and the edges that connect them reflect the P-values of their association. Network structure was visualized using the open-source software Gephi^45^. Networks were generated using different P-value thresholds (i.e., an edge was assigned only if the P-value was lower than 0.1,0.05 or 0.01). To determine robustness of network structure, we randomly deleted 10, 20 or 30 percent of all edges for each of the networks, and network topological properties were computed using the Cytoscape Network Analyzer tool^46^. Two metrics were calculated for the complete and depleted networks: **(i)** Degree distribution, and **(ii)** Closeness centrality^23^.

### Calculations of positive selection

We estimated for each codon of spike the number of inferred synonymous (S) and nonsynonymous (N) substitutions using the GALAXY platform ^47^. The input phylogenetic tree was constructed using FASTTREE. The dN-dS metric was used to detect codons under positive selection, where dS is the number of synonymous substitutions per site and dN is the number of nonsynonymous substitutions per site. dN-dS values were normalized using the expected number of substitutions per site. Maximum Likelihood computations of dN and dS were conducted using the HyPhy-SLAC software package^48^.

### Spatial clustering of volatility and calculations of the variable *D*

We performed a permutation test to determine the spatial clustering of volatile sites around each spike position. To this end, for each position *i*, we identified the 10 closest positions on the trimer, using coordinates of the cryo-EM structure of the cleavage-positive spike (PDB ID 6ZGI)^49^. We then calculated for each position *i* the statistic 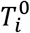:

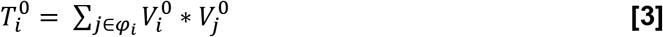

where 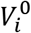 describes the volatility at position *i*, 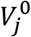 is the volatility at the j^th^ neighboring position to *i*, and *φ_i_* denotes the positions numbers of the 10 closest neighbors to position *i*. We then permuted all positions identifiers other than *p* and calculated the statistic 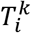:

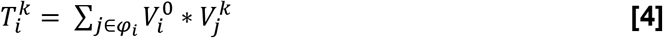

where 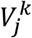 is the volatility at the j^th^ adjacent position in the *k*^th^ permutation (*k=1,2, … 5,000*).

Under the null hypothesis of no spatial clustering, we would expect the neighbor labels to be arbitrary. We therefore test this null hypothesis by estimating the probability of observing a positive departure from the null distribution via:

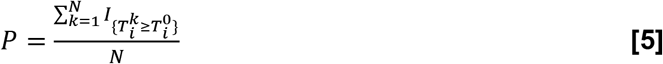

where *N* is the total number of permutations (5,000) and *I* is the indicator function. Therefore, the P-value quantifies the fraction of times the volatility of the surrounding residues is larger for the permuted values relative to the non-permuted values.

To calculate *D*, we measured for each position *i* the total volatility at all sites that are within a distance of 6 Å on the spike trimer structure. The coordinates of the cryo-electron microscopy structure of the cleaved spike protein in the closed conformation (PDB ID 6ZGI) were used^49^. Coordinates of all atoms were included; N-acetyl-glucosamine atoms were assigned the same position number as their associated Asn residues. We note that the 6ZGI structure is missing the following spike residues (numbered according to the SARS-CoV-2 reference strain): 1-13, 71-75, 618-632, 677-688, 941-943 and 1146-1273. To calculate *D* values for these positions, we applied the volatility values of the positions immediately adjacent on the linear sequence of spike (i.e., positions −1 and +1).

### Combined model to predict emergence of dominant-group and subgroup-emerging mutations

To assign a probability for each position to emerge with a mutation, we used a logistic regression model that applies *V, D* and *R* values. The model was trained using *V, D* and *R* values calculated using the 5,700 sequences of the baseline group, with the positive outcome being the 43 GDM and 16 sGEM sites described in **Supplementary Table 1**. To this end, we first created interaction terms between the initial predictors (i.e., *V, D* and *R*). To address the class imbalance in our datasets (59 of the 1,273 spike positions appeared with LFMs or sLFMs) we used the adaptive synthetic sampling approach (ADASYN)^50^. Nested cross-validation was used to tune the model while estimating the metrics of interest. This procedure was also used to generate the prediction probabilities for each position. Five folds were used for both the inner and outer parts of the nested cross-validation. Grid search was utilized to optimize hyperparameters with the area under the receiver operating characteristic curve as the objective for optimization. The model-specific parameters that we incorporated into the hyperparameter tuning procedure are the inverse of the regularization strength *C* and the penalty type. For this purpose, we used a set of values from 0.001 to 100 for parameter *C,* and for penalization we used L1 norm, L2 norm, elastic net, or no penalty in the parameter space. Since we used ADASYN to handle the class imbalance, we also added the number of positions with similar feature values as another hyperparameter to the search grid. The number of positions with similar feature values was set between 5 and 45. As classification metrics, we used sensitivity, specificity, precision, recall, AUC and balanced accuracy. The balanced accuracy metric, which is the average of sensitivity and specificity, was used due to the relative imbalance in the datasets.

## DATA AVAILABILITY STATEMENT

All IDs of the sequences used in our analyses are available on the Mendeley data repository at doi: 10.17632/wn7jwk9n22.1. Additional data related to this manuscript are available from the corresponding author upon request.

The custom code used in our studies is publicly available within the following hub repository: https://github.com/RoberthAnthonyRojasChavez/SARS2-Volatility. Source code for calculating lineage specificity can be found at https://github.com/haimlab/HIV.

## ACKNOWLEDGEMENTS

We are grateful to Dr. Wendy Maury and Dr. Stanley Perlman for critical reading of this manuscript. We are also grateful to Dr. Benjamin Darbro for helpful discussions. We extend our thanks to the GISAID consortium, NCBI Virus, ViPR and all laboratories that publicly summited their SARS-Cov-2 sequences. This work was supported by intramural funds to HH, by grant 110028-67-RGRL to HH from the American Foundation for AIDS Research (amfAR), and by National Institutes of Health grant 1DP2AI164325 to JD.

---

**Supplementary Figure 1.**
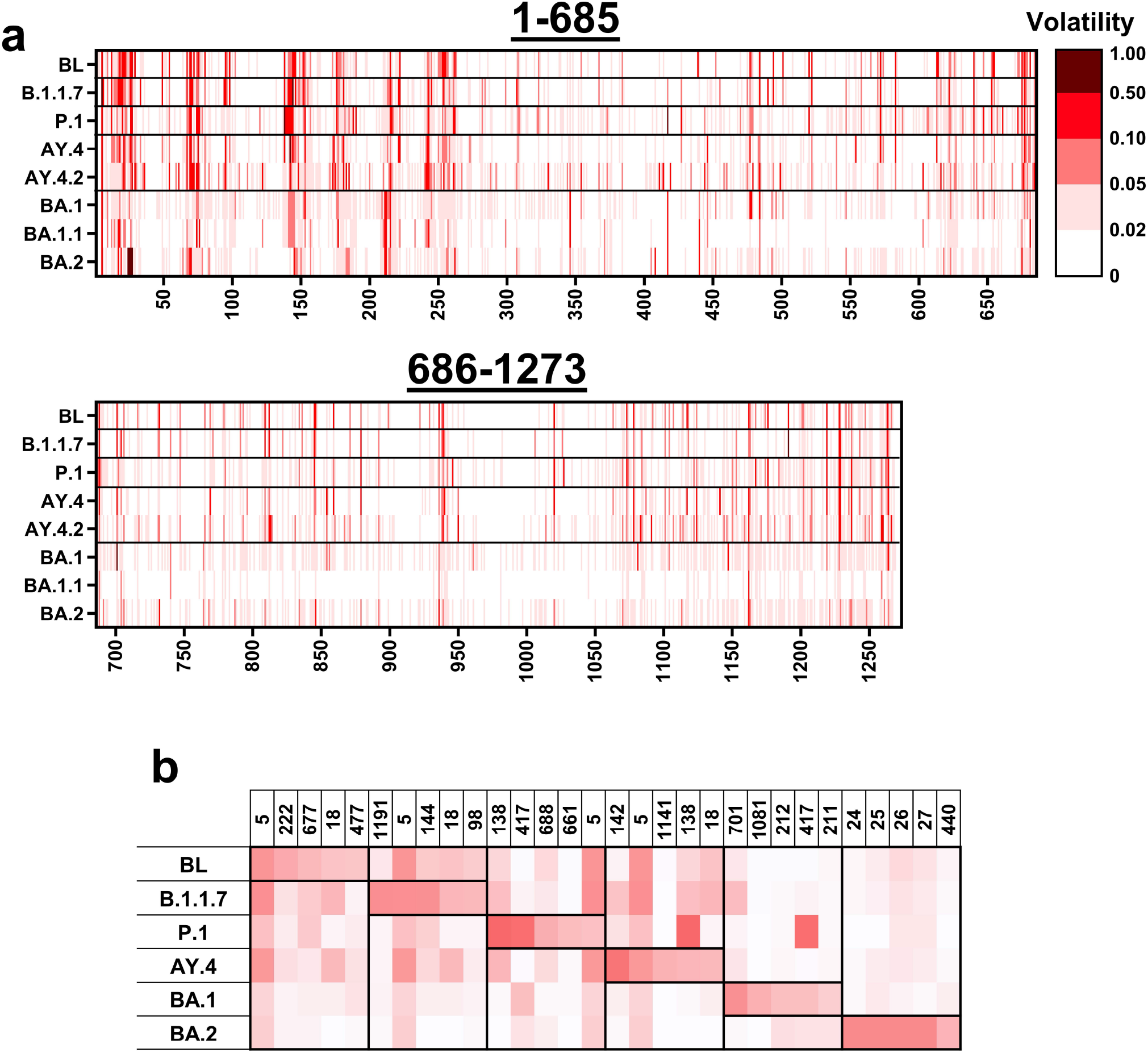
**(a)** Volatility calculated for all positions of spike using sequences from the indicated lineages or the SARS-CoV-2 baseline (BL) group. Data represent sequences from samples collected worldwide until the following dates: Baseline, until July 2021; P.1, AY.4, AY.4.2 and BA.1, until February 2022; B.1.1.7, BA.1.1 and BA.2, until March 2022. Volatility values are color-coded as indicated, **(b)** Volatilities are compared between lineages for the five positions with the highest values in each lineage.

**Supplementary Figure 2.**
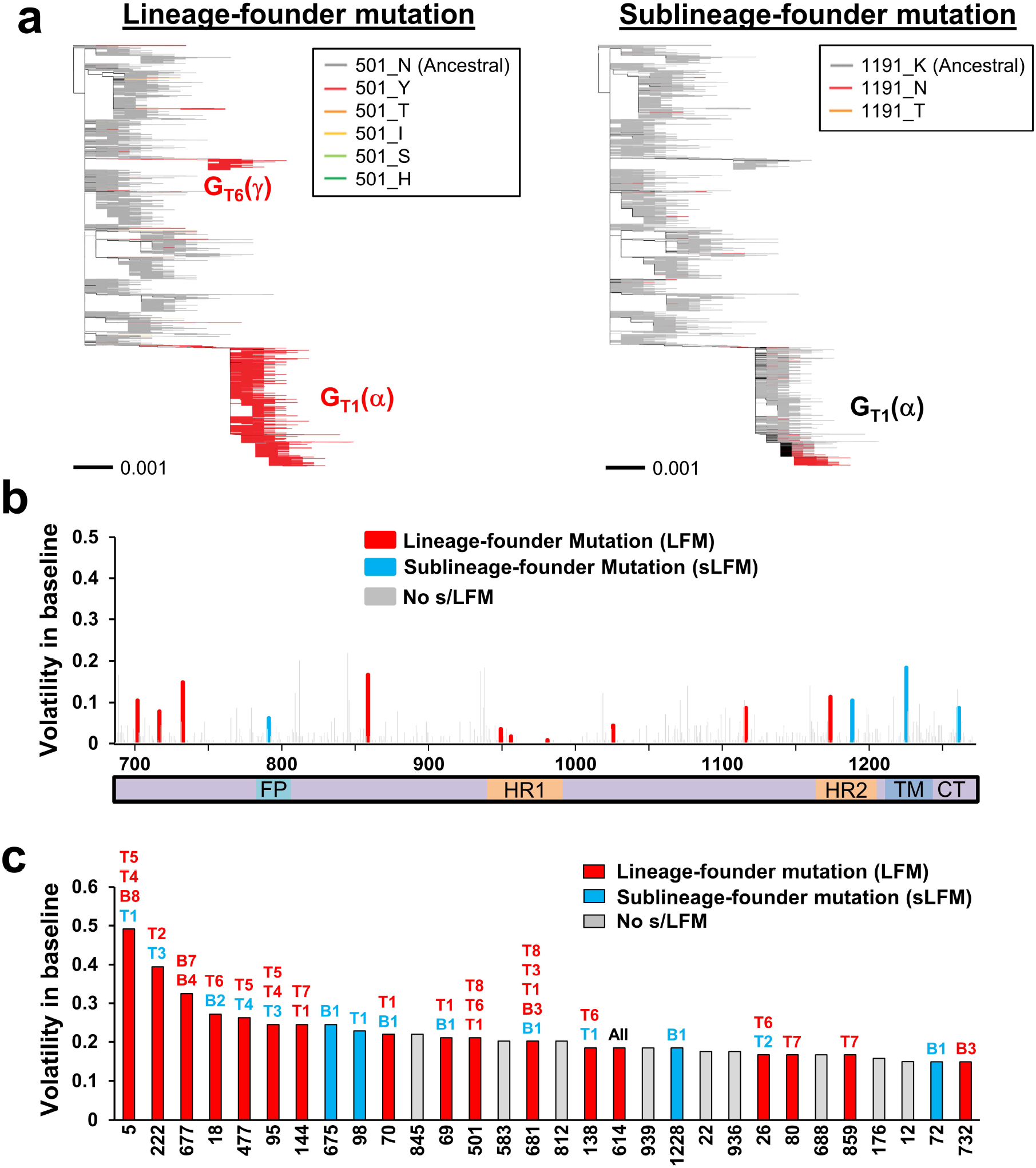
A high level of volatility at spike positions is associated with emergence of founder mutations in SARS-CoV-2 lineages. **(a)** Phylogenetic tree constructed from 16,808 unique spike sequences from samples collected worldwide until July 2021. Examples are shown of lineage-founder and sublineage-founder mutations in spike. (Left) Branches are colored by the amino acids that occupy spike position 501. The pattern corresponds to presence of a group-dominant mutation in G_T1_(α), G_T6_(γ) and G_T8_. (Right) Branches are colored by the amino acids that occupy position 1191, showing a sublineage-emerging mutation in G_T1_(α). **(b)** Volatility of spike positions of the S2 subunit, as calculated using the baseline group of 5,700 sequences (114 clusters). Red bars indicate positions with lineage-founder mutations (in any terminal or baseline group). Blue bars indicate positions with sublineage-founder mutations. FP, fusion peptide; HR, heptad repeats; TM, transmembrane domain; CT, cytoplasmic tail, **(c)** Thirty spike positions with the highest volatility values. The baseline (“B”) or terminal (“T”) groups that contain mutations at these positions are indicated.

**Supplementary Figure 3.**
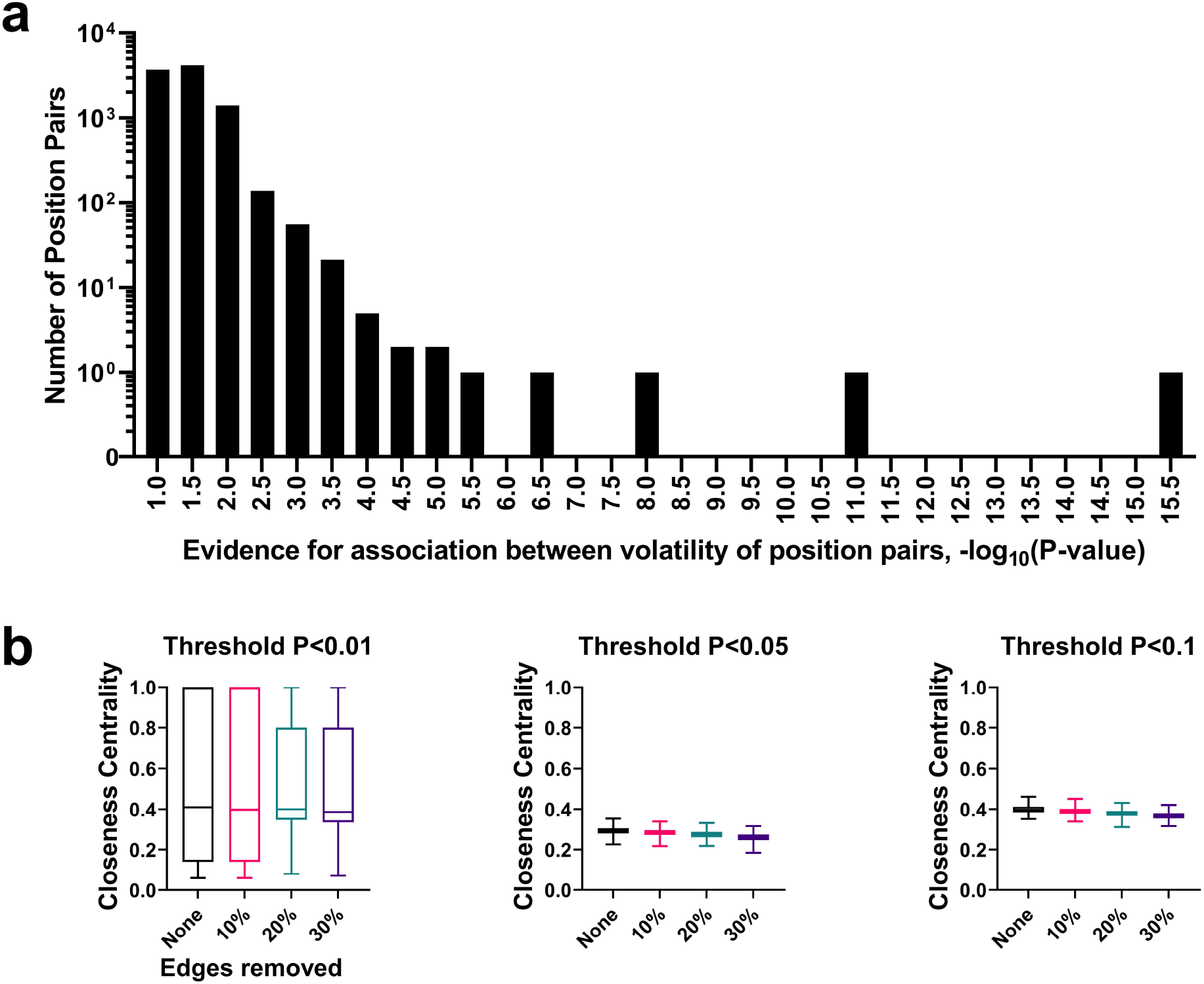
Structural properties of the network of co-volatile sites. The co-occurrence of a volatile state at any two spike positions was determined using the 114 clusters of the baseline group, **(a)** Histogram showing the distribution of P-values calculated using Fisher’s exact test for co-volatility of any two spike positions, **(b)** Networks of co-volatility between all spike positions were constructed using P-value thresholds of <0.01, <0.05 or <0.1. For each of the three networks, we randomly deleted 10%, 20% or 30% of edges and examined the effect on network stabilities. Closeness centrality values are shown for the intact and depleted networks. Higher values indicate shorter distances to all other nodes. Bars indicate the second and third quartiles and whiskers indicate minimum and maximum values.

**Supplementary Figure 4.**
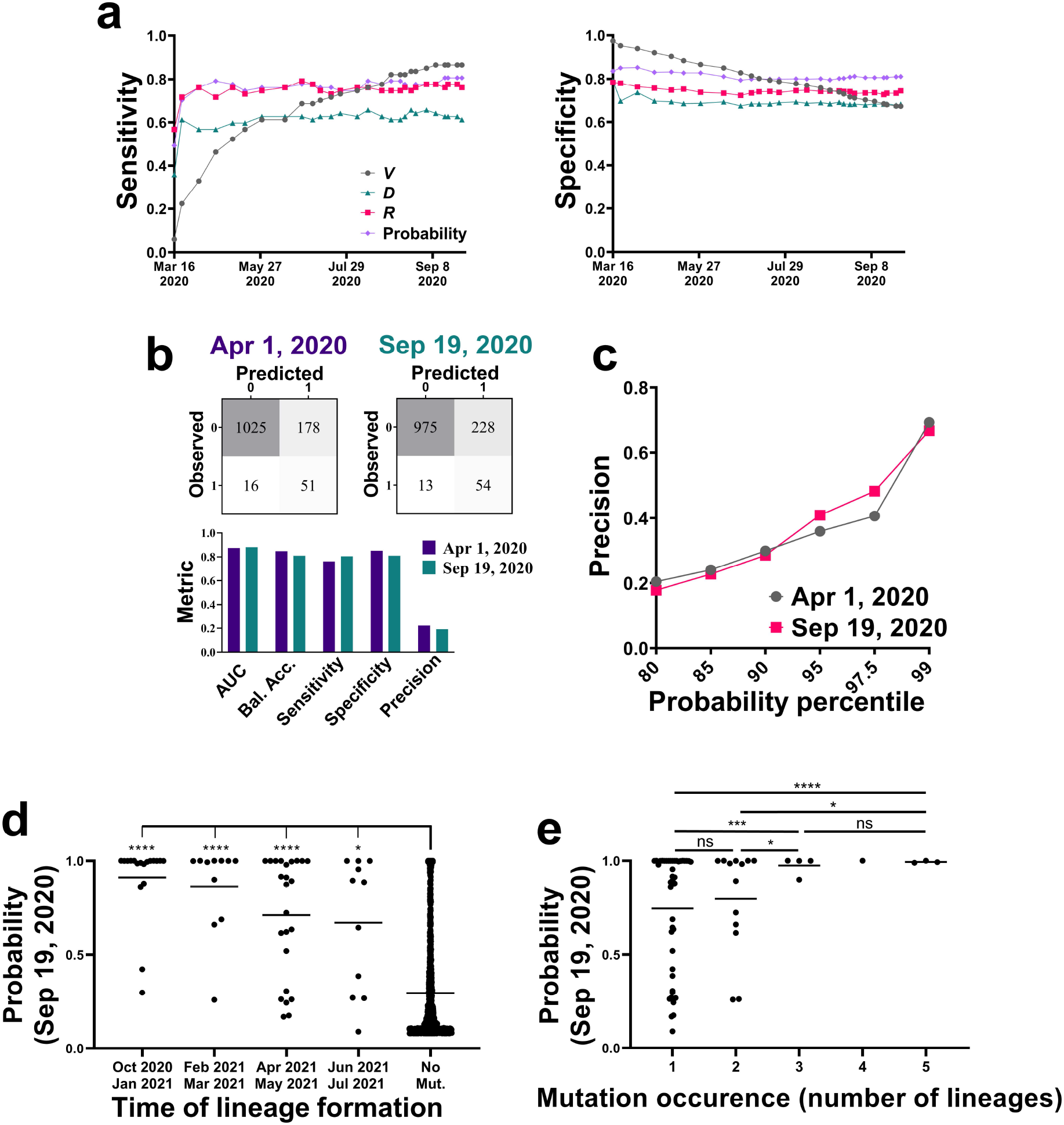
Volatility profiles among isolates from the early pandemic predict emergence of lineage-founder mutations at later stages. **(a)** Sequences from samples collected until the indicated time points of the early phase were used to calculate *V, D* and *R* values for each position. In addition, we calculated the probability for mutations at each spike position using a logistic regression model that applies *V, D* and *R* values. Sensitivity and specificity values are shown for prediction of the 67 LFMs that appeared during the lineage-emerging phase (September 2020 to July 2021). AUC values are shown in **Fig. 4f. (b)** Classification metrics for evaluating performance of the combined model to predict emergence of LFMs using sequence data collected until April 1^st^ 2020 or all sequences of the early phase, **(c)** Precision of the combined model at different probability percentile thresholds. Probabilities were calculated for all spike positions using the April 1^st^ 2020 dataset or all early-phase sequences, **(d)** LFM sites are grouped by the emergence time of the first lineage that contains them. Mutation probabilities assigned to the sites using sequences collected until September 19^th^ 2020 are shown and compared with the probabilities assigned to the no-mutation sites, **(e)** Comparison of probabilities assigned to LFM sites that appeared in one or more lineages. Probabilities assigned to positions in each group are compared between all groups using an unpaired T test: *, P<0.05, ***, P<0.0005; ns, not significant.

**Supplementary Figure 5.**
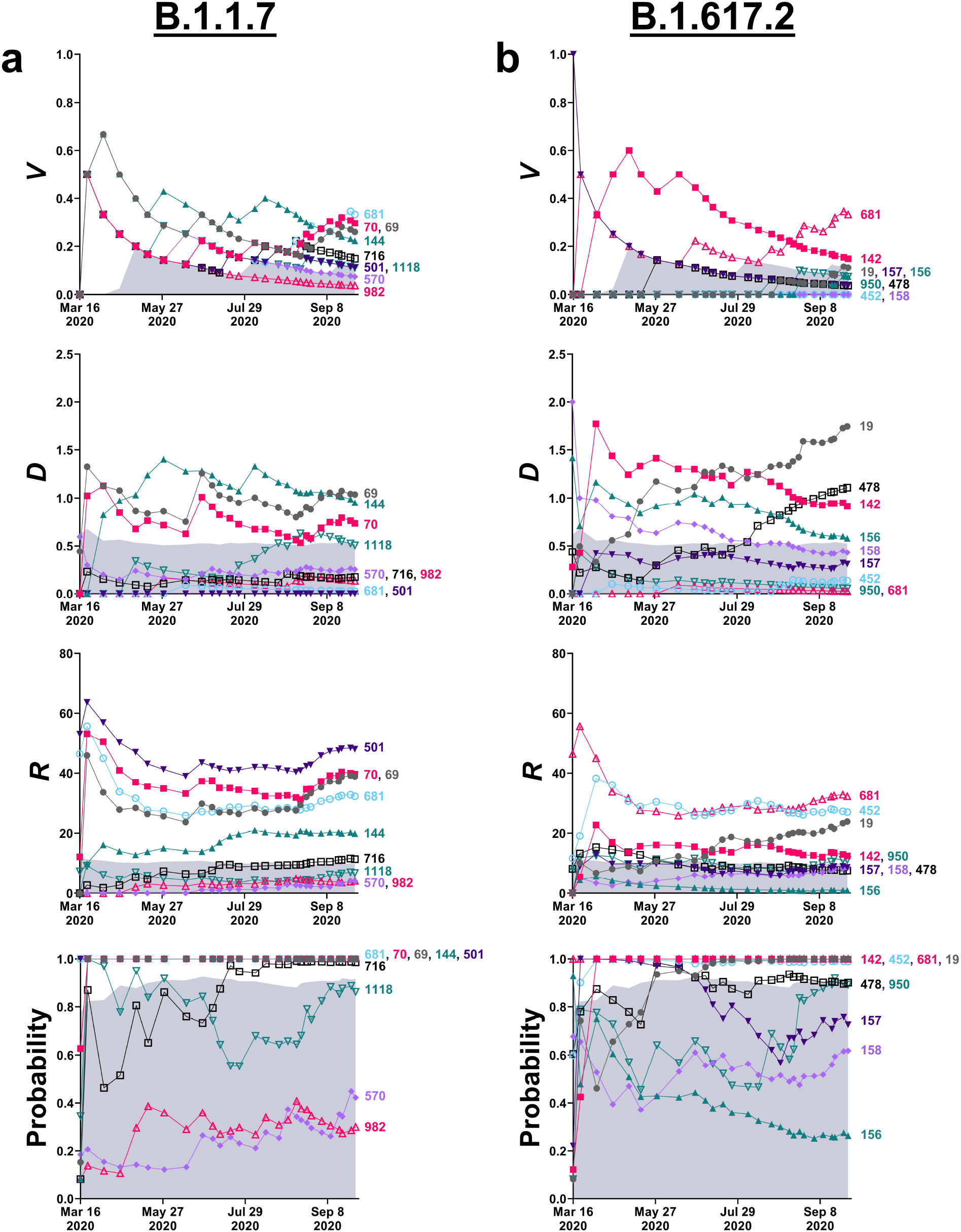
Evolution of *V, D* and *R* values and the combined mutation probability during the early phase of the pandemic. **(a,b)** Sequences from the early phase (December 30^th^ 2019 to September 19^th^ 2020) were divided into 36 clusters of 50 sequences. *V, D* and *R* values were calculated using sequences isolated from samples collected until the indicated time points. In addition, we calculated the probability for emergence of a mutation at each spike position using a logistic regression model that applies the three variables. Values for positions that emerged as LFMs in lineages B.1.1.7 and B.1.617.2 are shown. The shaded area describes probability values below the 90^th^ percentile calculated for each time point.

**Supplementary Table 1:**
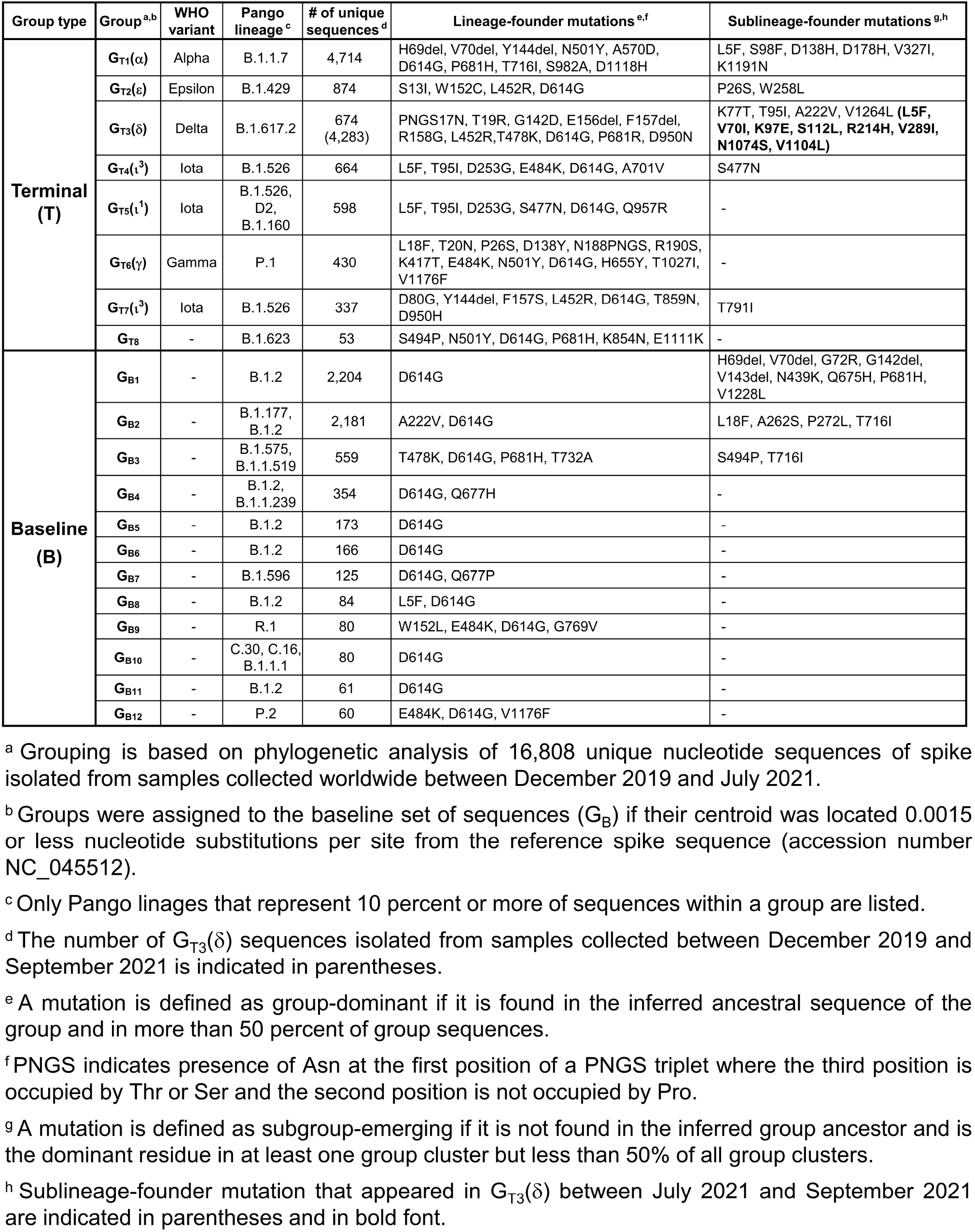
Founder mutations in SARS-CoV-2 lineages and sublineages.

**Supplementary Table 2:**
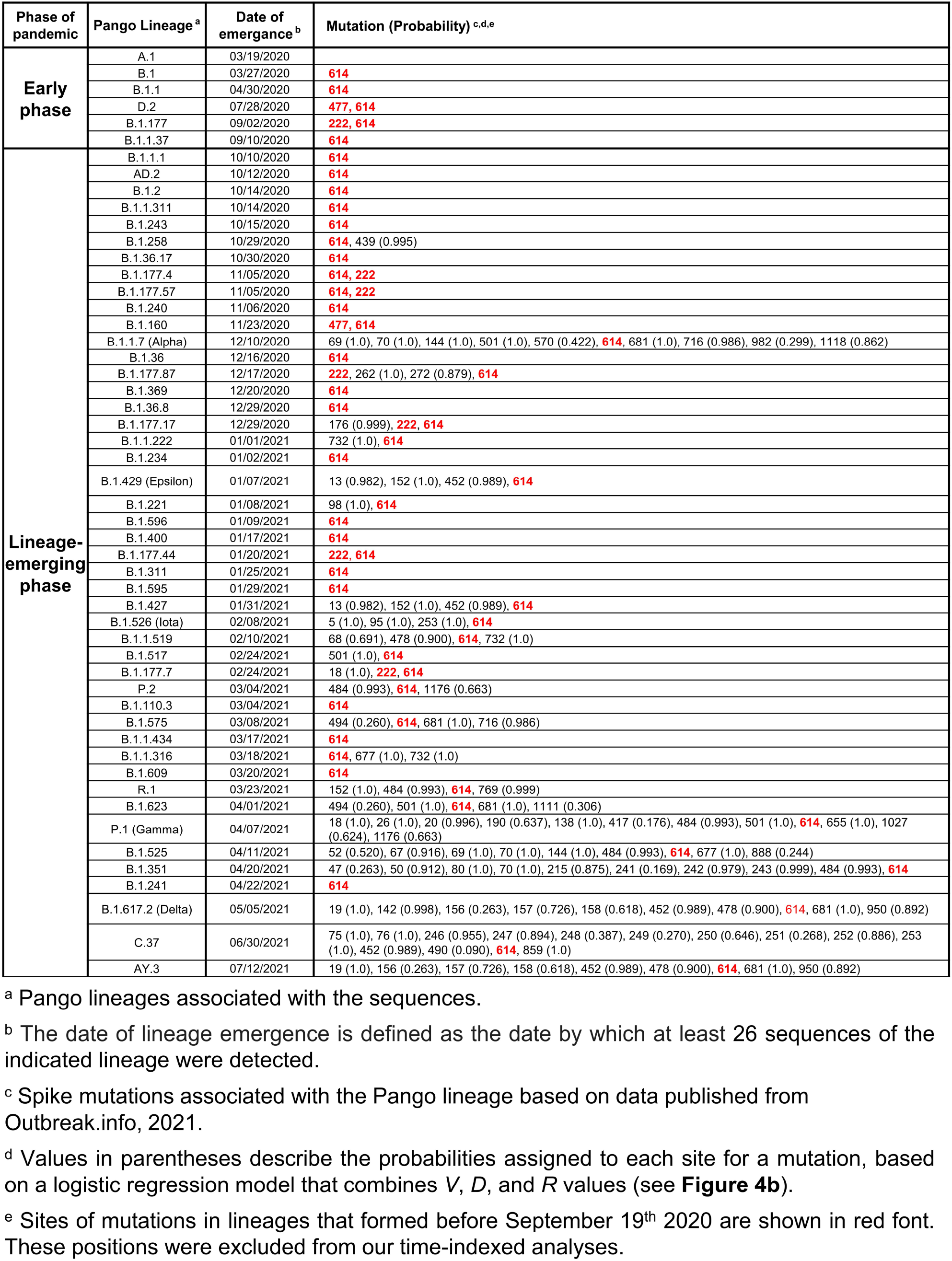
SARS-CoV-2 lineages that emerged until July 2021 and the mutations they contain in the spike protein.

| Phase of pandemic | Pango Lineage<sup>a</sup> | Date of emergence<sup>b</sup> | Mutation (Probability)<sup>c,d,e</sup> |
| --- | --- | --- | --- |
| Early phase | A.1 | 03/19/2020 |  |
|  | B.1 | 03/27/2020 | **614** |
|  | B.1.1 | 04/30/2020 | **614** |
|  | D.2 | 07/28/2020 | **477, 614** |
|  | B.1.177 | 09/02/2020 | **222, 614** |
|  | B.1.1.37 | 09/10/2020 | **614** |
| Lineage-emerging phase | B.1.1.1 | 10/10/2020 | **614** |
|  | AD.2 | 10/12/2020 | **614** |
|  | B.1.2 | 10/14/2020 | **614** |
|  | B.1.1.311 | 10/14/2020 | **614** |
|  | B.1.243 | 10/15/2020 | **614** |
|  | B.1.258 | 10/29/2020 | **614**, 439 (0.995) |
|  | B.1.36.17 | 10/30/2020 | **614** |
|  | B.1.177.4 | 11/05/2020 | **614, 222** |
|  | B.1.177.57 | 11/05/2020 | **614, 222** |
|  | B.1.240 | 11/06/2020 | **614** |
|  | B.1.160 | 11/23/2020 | **477, 614** |
|  | B.1.1.7 (Alpha) | 12/10/2020 | 69 (1.0), 70 (1.0), 144 (1.0), 501 (1.0), 570 (0.422), **614**, 681 (1.0), 716 (0.986), 982 (0.299), 1118 (0.862) |
|  | B.1.36 | 12/16/2020 | **614** |
|  | B.1.177.87 | 12/17/2020 | **222**, 262 (1.0), 272 (0.879), **614** |
|  | B.1.369 | 12/20/2020 | **614** |
|  | B.1.36.8 | 12/29/2020 | **614** |
|  | B.1.177.17 | 12/29/2020 | 176 (0.999), **222, 614** |
|  | B.1.1.222 | 01/01/2021 | 732 (1.0), **614** |
|  | B.1.234 | 01/02/2021 | **614** |
|  | B.1.429 (Epsilon) | 01/07/2021 | 13 (0.982), 152 (1.0), 452 (0.989), **614** |
|  | B.1.221 | 01/08/2021 | 98 (1.0), **614** |
|  | B.1.596 | 01/09/2021 | **614** |
|  | B.1.400 | 01/17/2021 | **614** |
|  | B.1.177.44 | 01/20/2021 | **222, 614** |
|  | B.1.311 | 01/25/2021 | **614** |
|  | B.1.595 | 01/29/2021 | **614** |
|  | B.1.427 | 01/31/2021 | 13 (0.982), 152 (1.0), 452 (0.989), **614** |
|  | B.1.526 (Iota) | 02/08/2021 | 5 (1.0), 95 (1.0), 253 (1.0), **614** |
|  | B.1.1.519 | 02/10/2021 | 68 (0.691), 478 (0.900), **614**, 732 (1.0) |
|  | B.1.517 | 02/24/2021 | 501 (1.0), **614** |
|  | B.1.177.7 | 02/24/2021 | 18 (1.0), **222, 614** |
|  | P.2 | 03/04/2021 | 484 (0.993), **614**, 1176 (0.663) |
|  | B.1.110.3 | 03/04/2021 | **614** |
|  | B.1.575 | 03/08/2021 | 494 (0.260), **614**, 681 (1.0), 716 (0.986) |
|  | B.1.1.434 | 03/17/2021 | **614** |
|  | B.1.1.316 | 03/18/2021 | **614**, 677 (1.0), 732 (1.0) |
|  | B.1.609 | 03/20/2021 | **614** |
|  | R.1 | 03/23/2021 | 152 (1.0), 484 (0.993), **614**, 769 (0.999) |
|  | B.1.623 | 04/01/2021 | 494 (0.260), 501 (1.0), **614**, 681 (1.0), 1111 (0.306) |
|  | P.1 (Gamma) | 04/07/2021 | 18 (1.0), 26 (1.0), 20 (0.996), 190 (0.637), 138 (1.0), 417 (0.176), 484 (0.993), 501 (1.0), **614**, 655 (1.0), 1027 (0.624), 1176 (0.663) |
|  | B.1.525 | 04/11/2021 | 52 (0.520), 67 (0.916), 69 (1.0), 70 (1.0), 144 (1.0), 484 (0.993), **614**, 677 (1.0), 888 (0.244) |
|  | B.1.351 | 04/20/2021 | 47 (0.263), 50 (0.912), 80 (1.0), 70 (1.0), 215 (0.875), 241 (0.169), 242 (0.979), 243 (0.999), 484 (0.993), **614** |
|  | B.1.241 | 04/22/2021 | **614** |
|  | B.1.617.2 (Delta) | 05/05/2021 | 19 (1.0), 142 (0.998), 156 (0.263), 157 (0.726), 158 (0.618), 452 (0.989), 478 (0.900), **614**, 681 (1.0), 950 (0.892) |
|  | C.37 | 06/30/2021 | 75 (1.0), 76 (1.0), 246 (0.955), 247 (0.894), 248 (0.387), 249 (0.270), 250 (0.646), 251 (0.268), 252 (0.886), 253 (1.0), 452 (0.989), 490 (0.090), **614**, 859 (1.0) |
|  | AY.3 | 07/12/2021 | 19 (1.0), 156 (0.263), 157 (0.726), 158 (0.618), 452 (0.989), 478 (0.900), **614**, 681 (1.0), 950 (0.892) |

<sup>a</sup> Pango lineages associated with the sequences.
<sup>b</sup> The date of lineage emergence is defined as the date by which at least 26 sequences of the indicated lineage were detected.
<sup>c</sup> Spike mutations associated with the Pango lineage based on data published from Outbreak.info, 2021.
<sup>d</sup> Values in parentheses describe the probabilities assigned to each site for a mutation, based on a logistic regression model that combines *V*, *D*, and *R* values (see **Figure 4b**).
<sup>e</sup> Sites of mutations in lineages that formed before September 19<sup>th</sup> 2020 are shown in red font. These positions were excluded from our time-indexed analyses.

**Supplementary Table 3:**
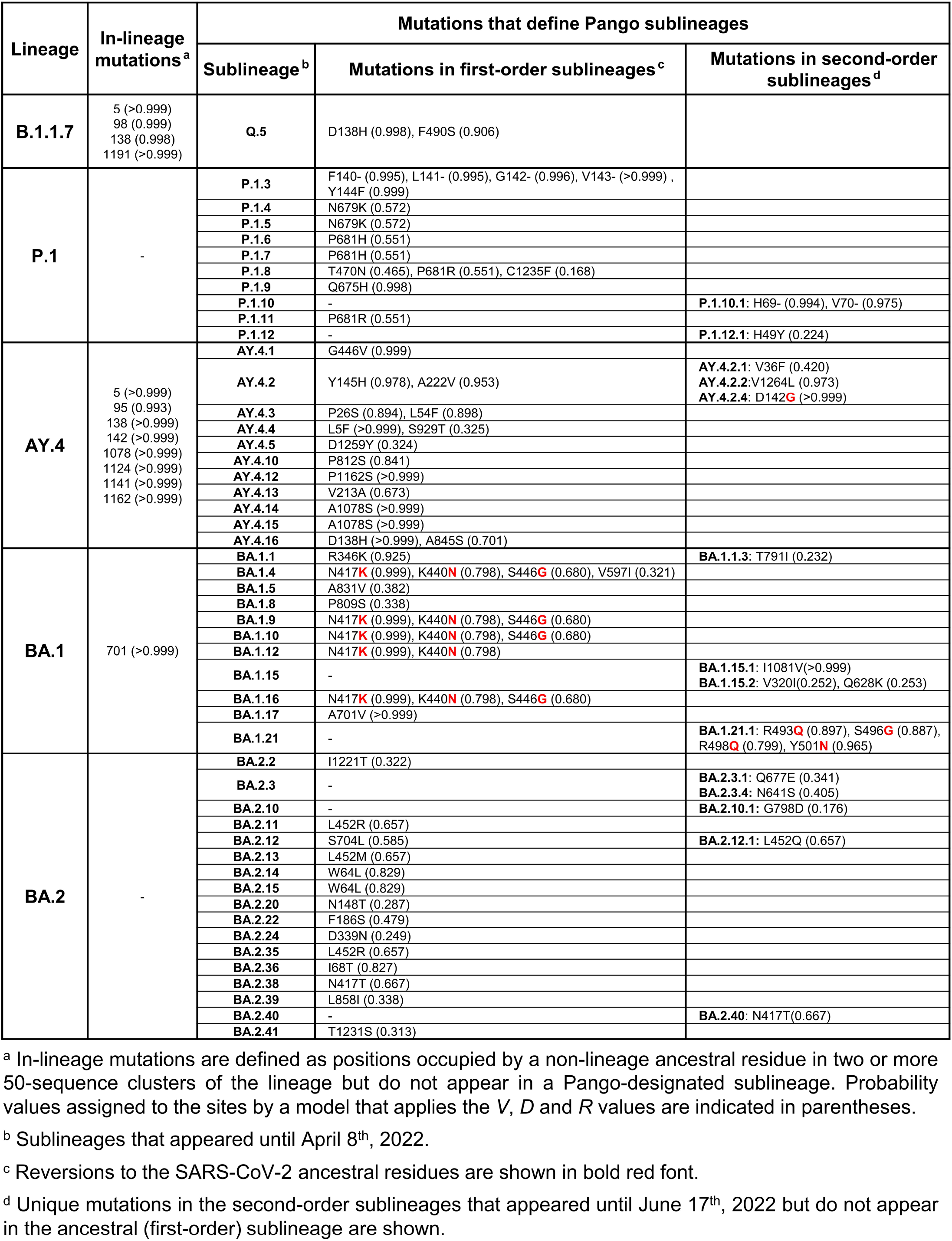
Mutations in spike that define SARS-CoV-2 sublineages.

